# Extension of individual model averaging assessments to unbalanced designs and dose-response

**DOI:** 10.1101/2024.04.13.589390

**Authors:** E. Chasseloup, X. Li, M. O. Karlsson

## Abstract

Recent investigations assessed two non-linear mixed effect (NLME) model based approaches to test for drug effect on real data in the context of balanced two-arms designs. The standard approach (STD) showed type I error inflation and biased drug effect estimates contrary to the proposed alternative, individual model averaging (IMA), which had controlled type I error and unbiased drug effect estimates. The current study is an extension of the performances assessment of these two approaches to unbalanced designs and dose-response studies. The type I error rate and drug effect estimates were assessed for unbalanced designs, using placebo Alzheimer disease assessment scale cognitive (ADAS-cog) scores from 800 individuals. The bias in the drug effect estimates was assessed for dose response scenarios, on data modified by the addition of various dose-response scenarios (Emax= 2.5, 5, and 10). The generalization of IMA to any randomization ratio of two-arms studies was also presented, together with an alternative parameterization of IMA: saturated IMA (sIMA). Similarly to what was observed in balanced designs, both IMA and sIMA had controlled type I errors and unbiased drug effect estimates in unbalanced designs, whereas STD had uncontrolled type I error and biased drug estimates. For the dose-response studies STD had a systematic bias towards the underestimation of the drug effect estimates. IMA and sIMA were unbiased in the scenarios with high maximum effect but their performances were hindered at the lowest maximum drug effect scenario, because of the closeness in magnitude between the drug effect addition and the placebo model misspecification.

## 2. Introduction

NLME models and “model-based drug development”[1] has been proven increasingly useful to support drug development and decision making[2–5]. Recent investigations assessed the performances (type I error and bias in drug effect estimates) of two NLME model-based approaches to test for drug effect in the context of two-arms balanced designs: STD and IMA[6]. Both approaches use the likelihood ratio test (LRT) to discriminate between a null hypothesis (*H*_0_), i.e. a base model without the design intervention information, and the alternative hypothesis (*H*_1_), i.e. a full model accounting for the design intervention information. The design intervention information being either the treatment allocation when testing for drug effect, or the combination of the treatment allocation and the dose administered when assessing dose-response. In STD, *H*_0_ describes the observations without any specific parameter describing the studied treatment intervention, while *H*_1_ adds a drug effect model to quantify the effect of the studied treatment intervention based on the arm assignment. IMA uses mixture models to test for the intervention effect. Both *H*_0_ and *H*_1_ hypotheses evaluate the individual likelihood (*IL*_*i*_) of two submodels on all individuals: a model with and a model without intervention effect. The difference between *H*_0_ and *H*_1_ is introduced in the mixture proportion estimate (*P*_*pop,Sk*_) used to average the individual likelihood of the two submodels according to Eqn. 1[7].

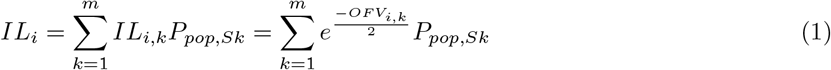

In the previous assessment of the two approaches, STD showed type I error inflation and biased estimates contrary to IMA which had controlled type I error and no biased estimates. The present work is an extension of the assessment of IMA and STD to more complex cases: unbalanced two-arms designs and dose-response. Real placebo data with or without the addition of simulated dose-response were used for the assessment of type I error and bias in drug estimates for STD, IMA and sIMA. The generalization of IMA to any randomization ratio of two-arms studies is detailed, together with an alternative parameterization of IMA called sIMA.

## 3. Methods

NONMEM version 7.4.4[8] with the FOCE estimation method without interaction was used for parameter estimation for all the models as the residual error model was additive. The randtest and sse (stochastic simulation and estimation) PsN[9] commands (version 4.9.0) automated the permutation tests and the fit of different models to data modified by the addition of simulated drug effect. The data management and post-processing of the results was carried out with R v3.6.2[10].

### 3.1 Original placebo data

The real placebo data were longitudinal ADAS-cog scores ranging from 0 to 70[11]. Due to the high number of categories the data were treated as continuous. 800 individuals were randomly selected for convenience among the recruited subjects (aged from 55 to 90 years old): 229 cognitively normal elderly, 188 presenting early Alzheimer disease, and 405 with mild cognitive impairment. The follow-up duration was respectively 3, 2, and 3 years, with ADAS-cog evaluation at 0, 6, 12, 24, and 36 months. The data set resulted in a total observation count of 3519. The Baseline Mini-Mental State (BMMS) was also collected at baseline for all the individuals and is used to describe the baseline ADAS-cog score.

### 3.2 Approaches

The three approaches described below (STD, IMA, and sIMA) discriminated between *H*_0_ and *H*_1_ with the LRT (*α* = 0.05). The LRT compared the likelihood ratio between *H*_0_ and *H*_1_ to the *χ*^2^ distribution, using the difference in parameters between the two hypotheses as degrees of freedom. NONMEM code examples for each approach are provided in the supplemental material 8.1.

#### 3.2.1 Standard approach (STD)

STD compared two nested models: *H*_0_, a function describing the evolution of the disease (*p*) without any treatment intervention model (Eqn. 2); and *H*_1_, which consisted of the same model with the addition of a function describing the treatment intervention (*d*) depending on the treatment allocation information (Eqn. 3).

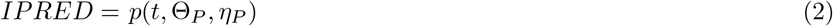

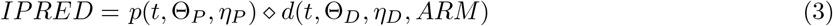

Where *IPRED* are the individual predictions, *t* is the time, Θ_*X*_ and *η*_*X*_ respectively the fixed and random effects of the functions, and ⋄ is an arithmetic operation (addition or multiplication). *ARM* ∈ { 0, 1} being the treatment allocation information (0: placebo arm, 1: treatment arm).

#### 3.2.2 Individual model averaging (IMA)

IMA discriminated between two nested models differing on the inclusion of the treatment allocation information. For both hypotheses a mixture model allowed the individuals to be allocated either to a submodel with a feature describing the treatment intervention (submodel *S*1 as described in Eqn. 3), either a sub-model without any treatment intervention feature (submodel *S*2 as described in Eqn. 2). While the *H*_0_ and *H*_1_ hypotheses contained each of the two submodels, they differed in the allocation probability to each of the submodels. In *H*_0_ the probability is fixed to the placebo allocation ratio (Eqn. 4), while in *H*_1_ this proportion is estimated (Θ_mix_) conditioned on the arm assignment (Eqn. 5).

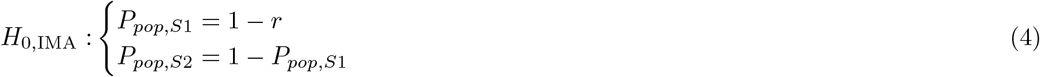

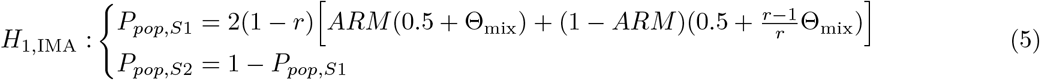

It is interesting to note that *P*_*pop,Sk*_ in *H*_0_ is equivalent to *P*_*pop,Sk*_ in *H*_1_ when fixing Θ_mix_ to 0. In order to keep the submodel probabilities within their definition domain (i.e., the interval [0, 1]) the following constraints are necessary when estimating Θ_mix_:

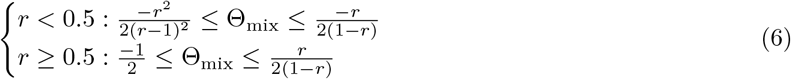

An illustration of the Θ_mix_ domain depending on the allocation ratio *r* is provided in supplemental material 8.2 in Figure 4.

#### 3.2.3 Saturated IMA (sIMA)

sIMA was an alternative implementation of IMA which used one additional parameter to estimate the submodel allocation probability in *H*_0_ (Eqn. 7), instead of fixing it to the placebo allocation ratio. In consequence, in *H*_1_ the probability of being allocated to each of the submodel was estimated through a different parameter for each ARM allocation (Eqn. 8). The final mixture proportion estimate of *H*_0_ was used as initial estimate for the two mixture proportions of *H*_1_.

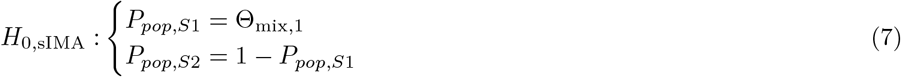

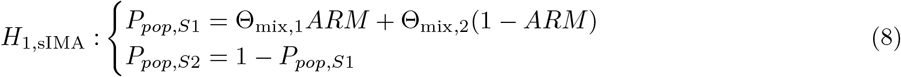

With Θ_mix,1_ ∈ [0, 1] and Θ_mix,2_ ∈ [0, 1]. To avoid the unblinding of the data caused by the usage of the dose-response model in *H*_0_ for the dose-response scenarios, the Emax model was replaced by an additional dose-independent offset term, called Extra. As a consequence, two alternatives were considered for *H*_1_ for the dose-response scenarios depending on whether the Extra term was kept in the treated submodel (see Table 1).

**Table 1:**
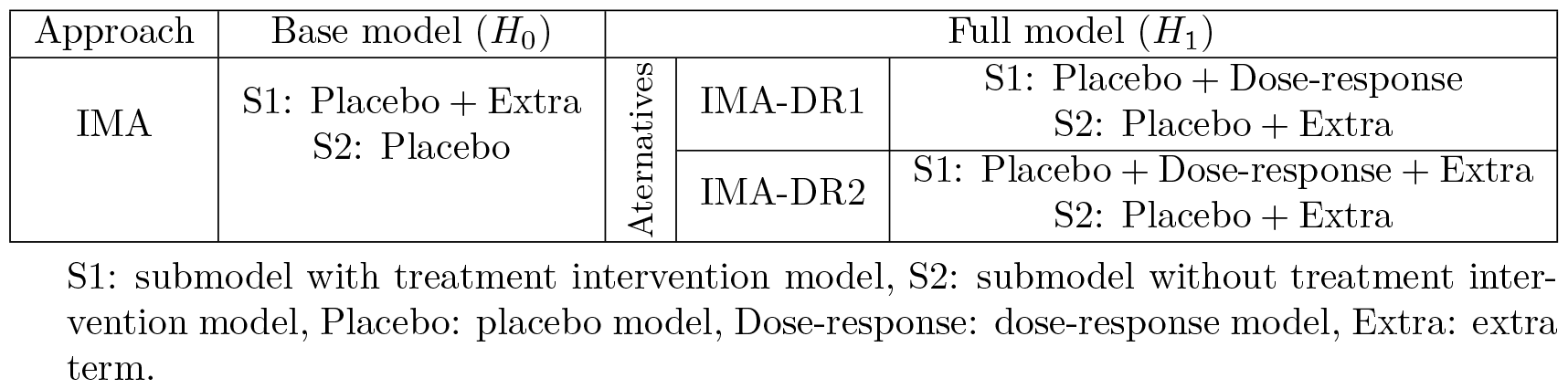
Details for the dose-response scenarios.

### 3.3 Type I error and bias in drug effect estimates assessment

The type I error was assessed in unbalanced studies for three different placebo allocation ratios (*r* = 0.25, *r* = 0.5, and *r* = 0.75), with 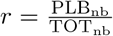, where PLB_nb_ is the number of individuals allocated to the placebo arm, and TOT_nb_ the sum of individuals allocated to both arms. Individuals were randomized repeatedly N=1000 times between the placebo arm (*ARM* = 0) and the treated arm (*ARM* = 1), to mimic N parallel group studies controlled with placebo and without drug effect. The type I error rate was computed as the frequency with which the approach concluded to a drug effect using the LRT. In STD the difference in parameters between *H*_0_ and *H*_1_ corresponded to the number of parameters of the drug model, while it is 1 for IMA and sIMA, corresponding to the estimation of the mixture proportion dependence on the treatment allocation. The nominal confidence interval for a type I error rate of 5 % was computed using a binomial distribution (*α* = 0.05, number of trials = N).

To simulate N randomized studies with dose-response, the individuals were randomized between placebo and treated arms using *r* = 0.25, and, when allocated to the treated arms, randomized a second time (1:1:1) between three different doses: 0.25, 0.667, and 4 (corresponding respectively to the dose required to achieve 20%, 40%, and 80% of the maximum drug effect). The time profiles were modified accordingly, with the addition of a dose response corresponding to an Emax model (dose-response model from Table 2). To evaluate the performances of the two approaches under various magnitudes of typical maximum drug effect, three scenarios with increasing Emax values were simulated (i.e. 2.5, 5 and 10), including 30% IIV on Emax.

**Table 2:** Details of the models used to assess the different approaches.

| Model | Name | Equation | Typical drug effect<sup>1</sup> |
| --- | --- | --- | --- |
| Placebo | linear | $\Theta_{\text{SLOPE}} \cdot t$ | - |
| Placebo | linear_iiv_slope | $(\Theta_{\text{SLOPE}} + \eta_{\text{SLOPE}})t$ | - |
| Placebo | exp_iiv_cor<sup>2</sup> | $(\Theta_{\text{Pmax}} + \eta_{\text{Pmax}}) \cdot (1 - e^{-\frac{\log(2)}{\Theta_{\text{HL}}} \cdot t})$ | - |
| Drug | offset | $\Theta_{\text{DRUG}}$ | $\Theta_{\text{DRUG}}$ |
| Drug | offset_iiv_drug | $\Theta_{\text{DRUG}} + \eta_{\text{DRUG}}$ | $\Theta_{\text{DRUG}}$ |
| Drug | linear | $\Theta_{\text{DRUG}} t$ | $\Theta_{\text{DRUG}} t_{\text{last}}$ |
| Drug | linear_iiv_drug | $(\Theta_{\text{DRUG}} + \eta_{\text{DRUG}})t$ | $\Theta_{\text{DRUG}} t_{\text{last}}$ |
| Dose-response | E<sub>max</sub> | $\frac{(\Theta_{\text{Emax}} + \eta_{\text{Emax}}) \text{Dose}}{\Theta_{\text{ED50}} + \text{Dose}}$ | $\frac{(\Theta_{\text{Emax}}) \text{Dose}}{\Theta_{\text{ED50}} + \text{Dose}}$ |
| Extra | Extra | $\Theta_{\text{EXTRA}}$ | $\Theta_{\text{EXTRA}}$ |

$\Theta$ are fixed effects and $\eta \sim \mathcal{N}(0, \omega)$ are random effects, the Pmax parameters describe the maximum placebo effect and the HL parameter the time required to reach half of this maximum effect. $t_{\text{last}}$ is the last observation time of the study, Dose=0.25, 0.667, 4, the Emax parameters describes the maximum drug effect and the ED50 parameter is the dose required to reach half of the maximum drug effect.
<sup>1</sup> Value computed at the end of the study. No typical drug effect for the placebo models. Corresponding to a typical drug effect when used as a drug model, or to a typical deviation when used as an extension of the placebo model.
<sup>2</sup> correlation between $\eta_{\text{Pmax}}$ and $\eta_{\text{Baseline}}$ .

The equation used for STD to compute the typical drug effect (TDE) of each drug model is detailed in Table 2. For IMA the typical deviation from the placebo trajectory and the relative estimated mixture proportion of each submodel, given the treatment allocation, was also accounted for in the computation of the TDE (Eqn. 9). For unbalanced designs, as the mixture describing the non-treated individuals consists only of the placebo model, *TD*_*S*2_ = 0, and the equation can be simplified to Eqn. 10.

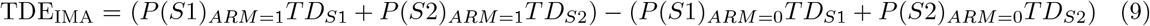

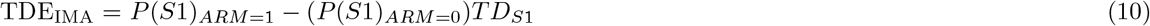

Where *P* (*SX*)_*ARM*=*Y*_ is the probability of being allocated to the submodel *X* for *ARM* = *Y*, and *TD*_*SX*_ is the typical deviation from the placebo trajectory for the submodel *X. TD*_*S*1_ = *f* (*Dose*) is evaluated at all the doses given in the corresponding arm, i.e. Dose = 0.25, 0.667, or 4 for *ARM* = 1, and Dose=0 hence *TD*_*S*1_ = 0 for *ARM* = 0, and *TD*_*S*2_ is the typical deviation quantified by the Extra term in *H*_1_. Eqn. 11 and 12 show the corresponding expressions of Eqn. 10 for IMA and sIMA respectively, when replacing the *P* (*SX*)_*ARM*=*Y*_ terms with the simplified expressions from Eqns. 4 and 8. Eqn. 13 describes the TDE for IMA-DR1, for which the Eqn. 9 cannot be simplified further.

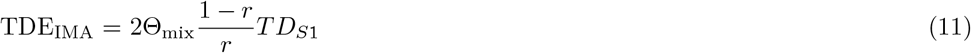

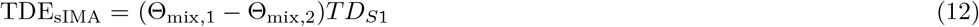

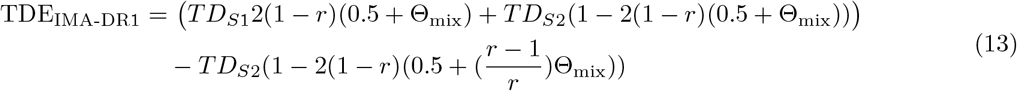

### 3.4 Models

The observed ADAS-cog scores of the individual i (*y*_*i*_) were described originally using Eqn. 14[11].

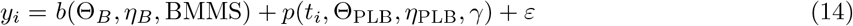

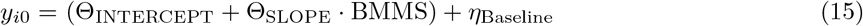

Where *b* is the function describing the observations at baseline (Eqn. 15). *t*_*i*_ is the individual time vector, Θ_*X*_ and *η*_*X*_ respectively the vectors of fixed and random effects of the functions, *γ* is the covariates vector, and *ε ∼ 𝒩* (0, *σ*).

For the unbalanced designs, two time-linear placebo models were tried, with and without IIV on the slope, and one exponential model with correlation between the maximum placebo effect and baseline. These three placebo models were combined with four different drug models: offset and time-linear with and without IIV. For the dose-response design the placebo model was the exponential model, and the dose-response model (Emax) used to simulate the dose-response scenarios. The equations for all the models are provided in Table 2.

## 4. Results

The type I error results for the three approaches, across the twelve placebo-drug models combinations are presented in Figure 1. More details regarding the minimization status are available in the supplemental material 8.3. The associated bias in the drug effect estimates at the end of the study are presented in Figure 2. Both IMA and sIMA had controlled type I error (*α* = 0.05) for all the placebo-drug models combinations tried while STD showed type I error inflation for the three allocation ratios (r=0.75, 0.5 and 0.25) and for most of the placebo-drug models combinations. For STD the type I error was controlled for the three placebo allocation ratios only for the three following combinations: the placebo linear_iiv_slope model combined with each of the two drug model without IIV, and the placebo exp_iiv_cor placebo model combined with the linear drug model.

**Figure 1:**
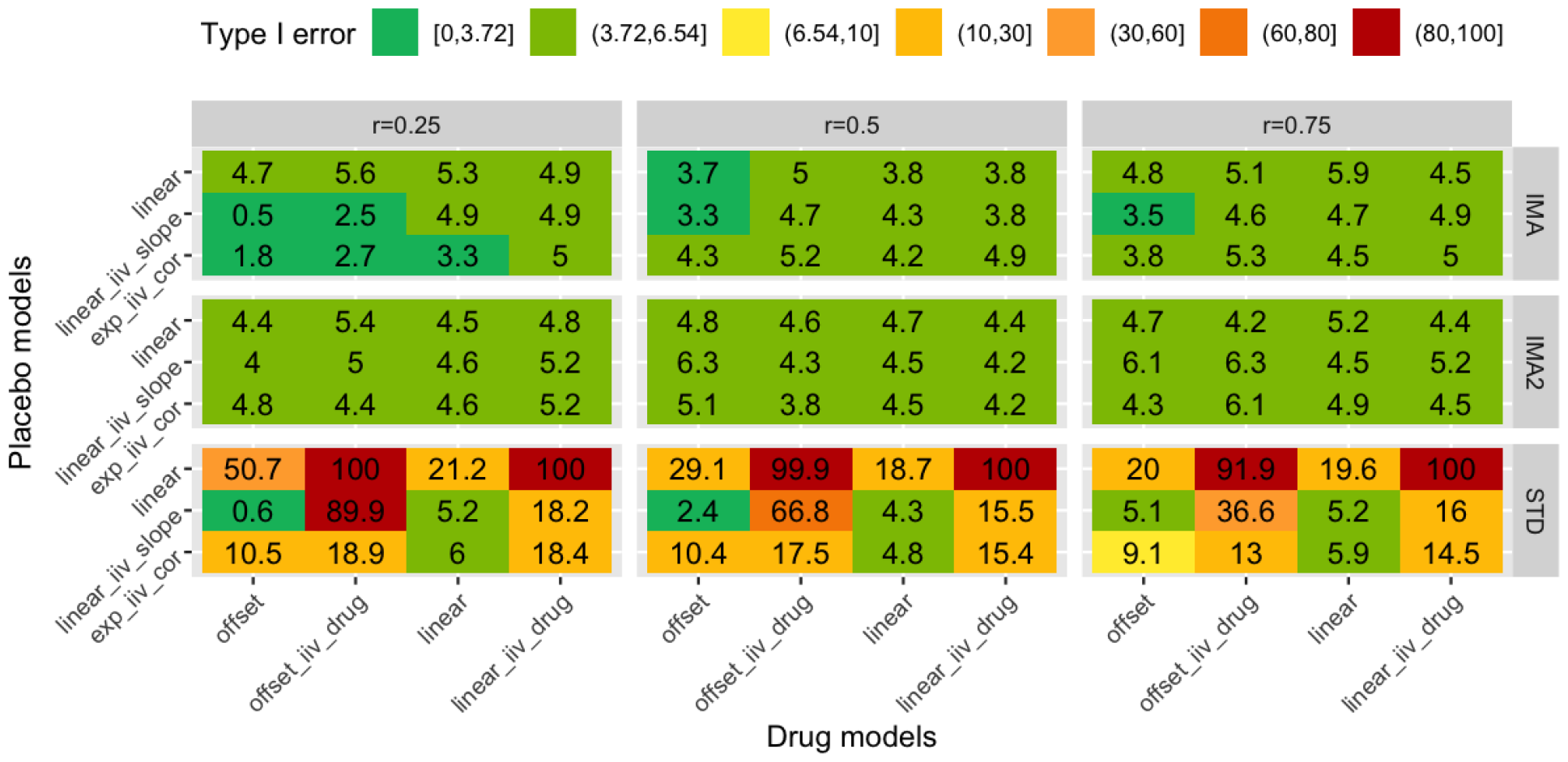
Type I error rate of STD, IMA, and sIMA for unbalanced designs using real placebo data (N=1000). Only the 3.72-6.54 type I error interval corresponds to a statistically relevant confidence interval. The other intervals were defined arbitrarily to help with visualization. The plot is facetted horizontally by placebo allocation ratio (r), and vertically by approach.

**Figure 2:**
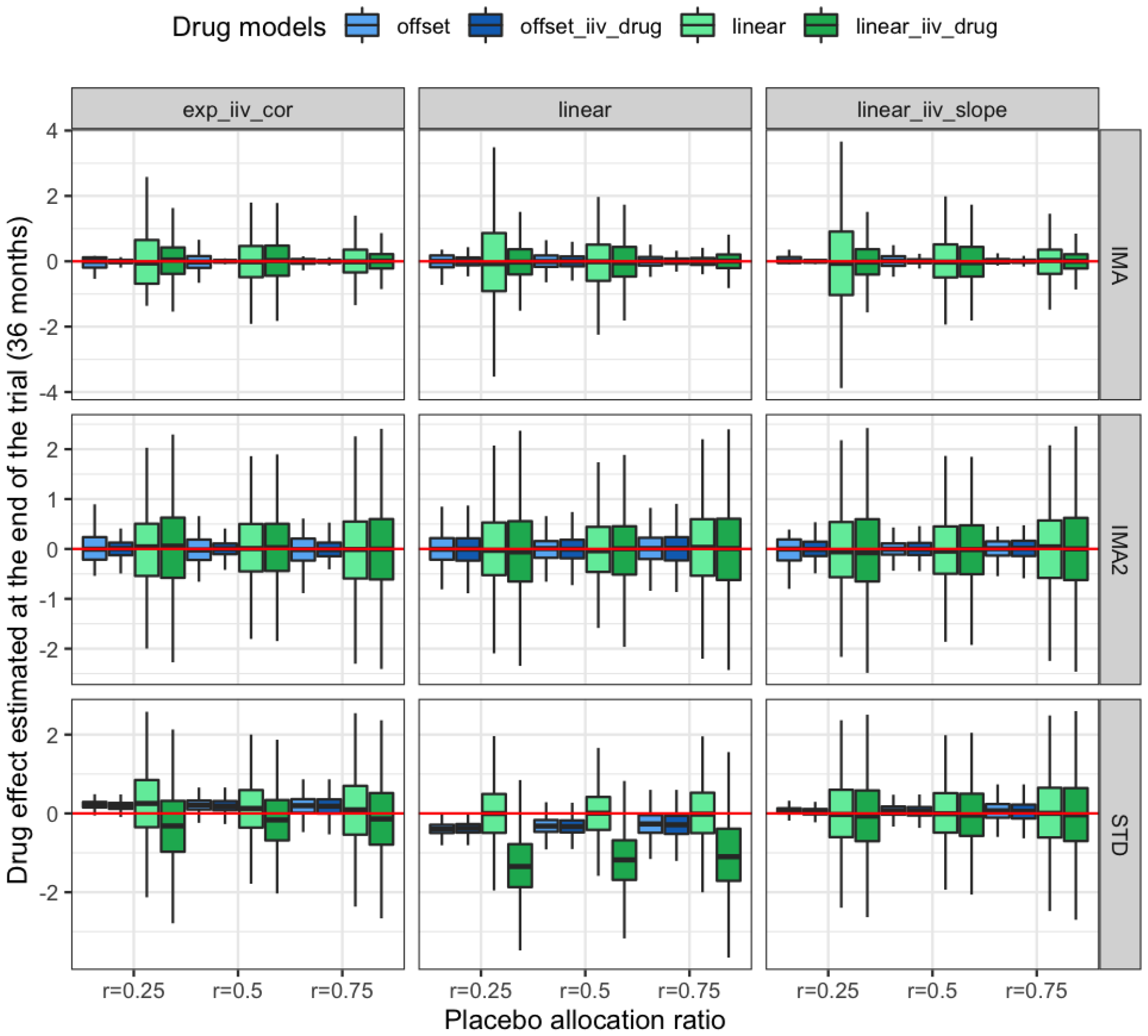
Bias in the drug effect estimated at the end of the trial (36 months) for STD, IMA, and sIMA for unbalanced designs using real placebo data (N=1000). The plot is facetted horizontally by placebo models and vertically by approach. The red line indicates the expected drug effect estimate of 0. r is the placebo allocation ratio.

Regarding the bias in the drug effect estimates, under the three placebo allocation ratios (r=0.75, 0.5 and 0.25), both IMA and sIMA had unbiased drug effect estimates. STD had randomly biased drug effect estimates, except for the linear drug models fitted with linear placebo models. The linear_iiv_slope placebo model was associated with the smallest biased drug estimates which were the highest for *r* = 0.25. Overall the offset drug models had more accurate estimates than the time-linear drug models. The placebo allocation had no major impact on the drug effect estimates, to the exception of IMA for which an increasing placebo allocation ratio increased the precision of the time-linear model drug estimates.

The results of the drug effect estimates for the dose-response scenarios are presented in Figure 3. Details regarding the minimization status are available in the supplemental material 8.4. STD systematically under-estimated the drug effect contrary to the two IMA implementations, with however a better accuracy. IMA had unbiased drug effect estimates for the simulated scenarios with a high maximum drug effect (Emax=5 and Emax=10). For the scenario with the lowest maximum drug effect (Emax=2.5), the two IMA approaches had better drug estimates than STD but were slightly underestimated especially for IMA-DR1.

**Figure 3:**
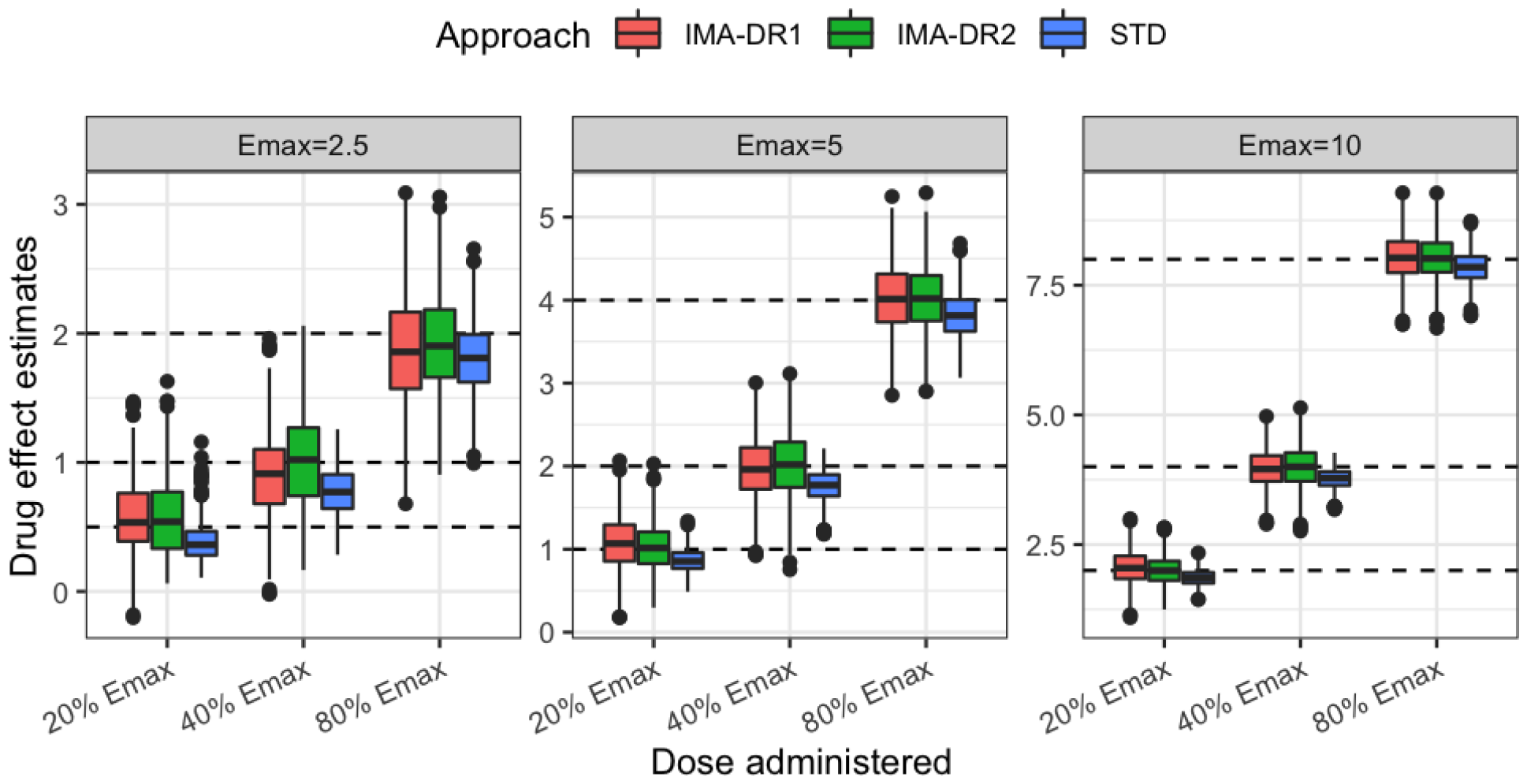
Drug effect estimates for each of the dose-response scenarios simulated (presented as horizontal facets), using the placebo data modified by the addition of a dose-response (N=1000). The dashed lines correspond to the values used in the simulations.

## 5. Discussion

This work extended the assessment of IMA and STD to the context of two-arms unbalanced designs and dose-response. Real placebo data were used to assess the type I error. The same data modified by the addition of a simulated dose-response were used to assess the bias in the drug effect estimates in such context.

The generalization of the equations to estimate the mixture proportion and to compute the drug effect with any placebo allocation ratio *r* was the first step to broaden the application of IMA further than two-arms balanced designs. It was a required step especially to assess IMA in the context of dose-response where multiple arms are treated with different doses. These general equations showed that the ones used previously[6] were a particular case where *r* = 0.5 and no typical deviation from the placebo model is estimated in the placebo submodel. Indeed, when *r* = 0.5, as in balanced designs, and Θ_mix_ = Θ_mix’_ *−*0.5, the *P*_*pop,S*1_ equation of *H*_1_ from Eqn. 5 is equivalent to *ARM* (1 *−*Θ_mix’_) + (1 *−ARM*)Θ_mix’_, the equation used previously. Similarly, the Eqn. 11 detailing the TDE computation for IMA is a generalization of (2Θ_mix’_ *−* 1)*TD*_*S*1_.

The results showed that IMA and sIMA had type I error control and unbiased drug effect estimates with unbalanced two-arms designs without drug effect. In the same context STD had uncontrolled type I error and randomly biased drug effect estimates. These results are in line with what was observed previously in the context of balanced two-arms designs[6]. In this work however an alternative implementation to IMA, sIMA, is introduced. Both allocate individuals to the most likely submodel using a mixture probability. In sIMA the mixture proportion is estimated in *H*_0_ instead of being fixed to a value dependent on the placebo allocation ratio, independently of the treatment allocation information. A different estimate is then allowed, conditioned on the treatment allocation in *H*_1_, using the final estimate from *H*_0_ to ensure that the models are nested which is a requirement to apply the LRT. As a result sIMA has an additional degree of freedom compared to IMA, both for *H*_0_ and *H*_1_.

When Θ_mix_ is estimated close to 0 in the full IMA model, or Θ_mix,1_ is estimated close to Θ_mix,2_ in the full sIMA model, the probability to be described by one the two submodels is the same for all the individuals, regardless of their arm allocation (*P* (*S*1)_*ARM*=1_ = *P* (*S*1)_*ARM*=0_). The full model is then similar to the the base model where *P* (*S*1) is not estimated based on the allocation information, which demonstrates that the treatment allocation is not a significant information to allocate the appropriate submodel to each individual. The estimated equiprobability based on the allocation information is a quantification of the absence of drug effect. An Θ_mix_ estimate in IMA, or Θ_mix,1_ and Θ_mix,2_ in sIMA, close to the limits of its domain indicates a strong correlation between the allocated submodel and the design treatment allocation (ARM). As these parameters represents probabilities, when the estimates are close to the boundaries, towards certainty, the allocation to the submodels becomes deterministic and is totally dependent on the ARM information. Inversely, given that *S*1 contains the drug model component, Θ_mix_ estimated equal to or near the lower boundary in the full IMA model, or Θ_mix,1_ estimated at the lower boundary of 0 and Θ_mix,2_ at the upper boundary of 1 in the full sIMA model (i.e., *P* (*S*1)_*ARM*=1_ = 0 and *P* (*S*1)_*ARM*=0_ = 1) would suggest a model misspecification.

Both IMA and sIMA had a type I error control, however as the placebo allocation was decreasing, IMA had increasing number of placebo-drug models combinations with a type I error rate that was even lower than the binomial confidence interval built around the 5% reference value with a p-value of 0.05. Only sIMA had type I error rate within the confidence interval across all the scenarios. These results suggest that sIMA could have a better power than IMA. The type I error for STD show that with this approach, the type I error was always inflated when the drug models had IIV, suggesting than the placebo model was largely misspecified in that respect. This also suggests that it could be more reasonable to test for drug effect with drug models describing only the typical tendency. Two additional arguments points in that direction: (1) the drug models with IIV had higher type I error rate when associated with the placebo models without IIV term (the linear placebo model); and (2) the drug models without IIV were associated with the lowest or controlled type I error rates. The type I error rate was also lower when the drug model was the same mathematical function as the placebo model, i.e. when the drug model did not offer a real additional degree of freedom. With the exp_iiv_cor model, the combination of the PMAX and HL parameters resulted in a linear behavior within the time frame of the study. This explains that the behavior of the exp_iiv_cor and the linear_iiv_slope placebo models were similar when associated with the linear drug models. The association of this placebo exp_iiv_cor model with the offset drug models resulted in higher type I error rate since these drug models were more dissimilar to the placebo model.

Regardless of the approach, offset drug models resulted in more accurate drug estimates at the end of the study than the time-linear drug models. This can be explained by the similarity between the placebo models and the drug effect models for the time-linear drug models. The precision is lower for sIMA than IMA but this is in line with the fact that an additional parameter was systematically estimated in sIMA, and consequently also included in the computation of the typical drug effect. For STD, the linear drug model which is the drug model that likely offered the less additional degrees of freedom, was the least biased. The linear_iiv_slope placebo model was the model associated with the least biased drug effect estimates, presumably because it was the least misspecified model as it is the closest to the published placebo model[11].

The *r* = 0.5 scenario correspond to the two-arms balanced design that was studied before[6], the results differ slightly however because the data were modified in that study by adding -1 to the whole time vector to allow the observations record to start at time=0 (the observations were registered from month=1 initially). This change allows for a time-linear drug effect of 0 at time 0 and explains why a better type I error rate was observed in this study for the combination of the placebo “linear_iiv_slope” model and the drug “linear” model.

Regarding the results for the placebo data modified by the addition of a simulated dose-response effect, the exp_iiv_cor model was picked as it was the placebo model with the lowest OFV. Both IMA-DR1 and IMA-DR2 showed unbiased drug effect estimates when the maximum drug effect was high enough. On the contrary STD systematically underestimated the drug effect, regardless of the magnitude of the maximum drug effect simulated, however with better precision. The Θ_mix_ parameter of both IMA-DR1 and IMA-DR2 approaches were estimated to the upper boundary, leading to a deterministic submodel allocation, entirely based on the ARM allocation. For the lowest maximum drug effect simulated (Emax = 2.5), IMA-DR1 was more biased, underestimating slightly the drug effect. IMA-DR2 was also underestimating slightly the drug effect at the highest dose of the lowest Emax scenario. A look at the mixture proportions estimates (Figure 8 in supplemental material 8.4) show that at the lowest Emax scenario, the median of the mixture proportion was lower for IMA-DR1. As the mixture proportion is included in the computation of the final drug estimates, values lower than the upper boundary tend to decrease the final estimates. An other information from this mixture proportion estimates is that not only the median is lower for IMA-DR1, but for both IMA approaches the distribution is different for the lowest Emax scenario. Both have a the inter-quartile distribution ranging from the lower bound to the higher bound of the proportion domain. This behavior showed that with this magnitude of Emax, the IMA approaches were not always assigning the individuals with a drug effect to the model with the dose-response function. One hypothesis for this difficulty it that the maximum effect addition with this scenario is of the same magnitude as the size of the model misspecification. It is interesting to note that as the formula to compute the drug effect for the IMA approaches accounts for the deviation from the typical placebo trajectory included in the placebo submodel (i.e. the Extra term in *S*2), the final drug estimate was still reasonable. Additional studies with other data set from other disease area would be necessary to decide whether this threshold is data dependent or approach dependent.

STD was both more precise but more biased than the IMA approaches, underestimating systematically the drug estimate across the three dose-response simulated scenarios. The main hypothesis for this observed difference in performances in terms of bias for is the absence in STD of the Extra term. In the IMA approaches, this typical deviation from the placebo trajectory estimated on the non-treated individuals is removed from the final drug estimates of the treated individuals. The remaining misspecification from the placebo model is then removed from the final drug estimate, consequently increasing the accuracy. In STD this potential model placebo misspecification is not estimated and hence not removed. However this additional parameter comes with a cost which decreases the estimates precision in the final estimates of the IMA approaches, in addition to the additional parameter brought by the mixture proportion estimate already mentioned before.

The Extra term was introduced in the base IMA models to allow the estimation of a drug effect without unblinding the data, which would have been inevitable when using the Emax model. This offset term allows a time independent description of the drug response without using neither the dose or the arm information. In the full IMA models, this Extra term can be interpreted as a time independent quantification of the placebo model misspecification.

Extensions to more placebo-drug models combinations, in different disease areas, and different drug effect or dose-response models would be interesting eventually to generalize the observations.

## 6. Conclusion

This study focused on the extensions of IMA and STD performances assessment using real data, to unbalanced designs for two-arm studies and dose-response. IMA was successfully generalized to any placebo allocation ratio and an additional implementation of IMA, called sIMA, was also proposed with similar properties. Contrary to IMA, sIMA allows the estimation of the submodel allocation proportion in the base model, to the cost of an additional parameter to estimate.

For unbalanced designs, the results were similar to the results observed previously with balanced designs. Both IMA and sIMA had controlled type I error and unbiased drug effect estimates, contrary to STD which had inflated type I error and randomly biased estimates on data without drug effect. Similar behavior was observed when using data modified by the addition of a simulated dose-response, leading to a systematic bias for STD, and no bias for IMA approaches, except in the lowest simulated Emax scenario where IMA could evidence the similarity in size of the magnitude of the drug effect estimates and the model misspecification.

The proposed explanation to the difference in performances seems confirmed in the unbalanced designs: in STD any feature of the data not described by the placebo models is likely to make a new model feature significant, leading to bias in drug effect estimates and inflated type I errors rates when drug models provide more flexibility/degrees of freedom/parameters to describe the data. IMA does not suffer from this problem since placebo and drug models are fitted together to the whole data set, both in base and full models, and was able to integrate in the final drug effect estimates a quantification of the placebo model misspecification.

## 8. Supplemental material

### 8.1 NONMEM code examples

#### 8.1.1 Standard base model

~~~
## $PROBLEM ADAS-cog
## $INPUT ID TIME DV BMMS ARM ;$INPUT C ID DROP TIME DV MMSE BADS BMMS DROP DROP DROP
## ;PSTS AGE SEX DROP EDU APOE APOF FHAD ABET DROP MDV INVF ARM
## ;ADAS=DV ##
## ;TIME in month ##
## ;BADS: baseline ADAScog
##
## ;BMMS: baseline MMSE
##
## ;;PSTS: 1=normal, 2=MCI, 2=AD
##
## ;APOF: ApoE 0=non carrier, 1=hetero, 2=homo-carrier
##
## ;FHAD: family history of AD, 1=either M or F is AD, 0=no, or NA
## $DATA ../cleaned_800_13_dur36_exp_iiv_0_omg0.csv IGNORE=@
## $ABB COMRES=3
## $PRED
##
## ;Baseline model
## IF (NEWIND.LE.0) THEN
##    LN2=LOG(2)
## ENDIF
##
## INTERCEPT = THETA(1)
## BSLP      = THETA(2)
## TVBSL     = INTERCEPT + BSLP * BMMS
## BASELINE  = TVBSL + ETA(1)
##
## ;----PLACEBO MODEL-----
## PMAX    = THETA(4)+ETA(2)
## HL      = THETA(5)
## PLACEBO = PMAX*(1-EXP(-LOG(2)/HL*TIME))
##
## ;----DRUG MODEL-----
## DRUG=0 ; drug_model
##
## ADASCOG = BASELINE + DRUG + PLACEBO
## IF(TIME.EQ.0) ADASCOG = BASELINE + PLACEBO
##
## F = ADASCOG
## ADD = THETA(3)
## W = ADD
## IPRED = F
## Y = F + W * EPS(1)
##
## $THETA (0,52.7449) ; PRM INTER 1
## $THETA -1.53777 ; PRM BSLP 2
## $THETA (0,2.81553) ; PRM EPS 3
## $THETA 21.2003 ; PRM PMAX 4
## $THETA (0,85.3397,100) ; PRM HL 5
##
## $OMEGA BLOCK(2)
## 15.3857 ; PRM OMBASE 1
## 70.2731 1277.18 ; PRM OMPMAX 2
##
## $SIGMA 1 FIXED
##
## $ESTIMATION MAXEVAL=9999 METHOD=1 NOABORT NOTHETABOUNDTEST NOOMEGABOUNDTEST
~~~

#### 8.1.2 Standard full model

~~~
## $PROBLEM ADAS-cog
## $INPUT ID TIME DV BMMS ARM ;$INPUT C ID DROP TIME DV MMSE BADS BMMS DROP DROP DROP
##
## ;PSTS AGE SEX DROP EDU APOE APOF FHAD ABET DROP MDV INVF ARM
##
## ;ADAS=DV
##
## ;TIME in month
##
## ;BADS: baseline ADAScog
##
## ;BMMS: baseline MMSE
##
## ;;PSTS: 1=normal, 2=MCI, 2=AD
##
## ;APOF: ApoE 0=non carrier, 1=hetero, 2=homo-carrier
##
## ;FHAD: family history of AD, 1=either M or F is AD, 0=no, or NA
## $DATA ../cleaned_800_13_dur36_exp_iiv_0_omg0.csv IGNORE=@
## $ABBREVIATED COMRES=3
## $PRED
##
## ;Baseline model
## IF (NEWIND.LE.0) THEN
##   LN2=LOG(2)
## ENDIF
##
## INTERCEPT = THETA(1)
## BSLP      = THETA(2)
## TVBSL     = INTERCEPT + BSLP * BMMS
## BASELINE  = TVBSL + ETA(1)
##
## ;----PLACEBO MODEL-----
## PMAX    = THETA(4)+ETA(2)
## HL      = THETA(5)
## PLACEBO = PMAX*(1-EXP(-LOG(2)/HL*TIME))
##
## ;----DRUG MODEL-----
## DRUG=THETA(6)*TIME ; drug_model
##
## ADASCOG = BASELINE + DRUG*ARM + PLACEBO
## IF(TIME.EQ.0) ADASCOG = BASELINE + PLACEBO
##
## F = ADASCOG
## ADD = THETA(3)
## W = ADD
## IPRED = F
## Y = F + W * EPS(1)
##
## $THETA (0,52.7449) ; PRM INTER 1
## $THETA -1.53777 ; PRM BSLP 2
## $THETA (0,2.81551) ; PRM EPS 3
## $THETA 21.2001 ; PRM PMAX 4
## $THETA (0,85.3398,100) ; PRM HL 5
## $THETA (1) ; PRM DRG 6
## $OMEGA BLOCK(2)
##         15.386 ; PRM OMBASE 1
##         70.2726 1277.17 ; PRM OMPMAX 2
## $SIGMA 1 FIX
## $ESTIMATION MAXEVAL=9999 METHOD=1 NOABORT NOTHETABOUNDTEST
##                    NOOMEGABOUNDTEST
~~~

#### 8.1.3 IMA base model

~~~
## $PROBLEM ADAS-cog
## $INPUT ID TIME DV BMMS ARM ;$INPUT C ID DROP TIME DV MMSE BADS BMMS DROP DROP DROP
## ;PSTS AGE SEX DROP EDU APOE APOF FHAD ABET DROP MDV INVF ARM
## ;ADAS=DV
##
## ;TIME in month
##
## ;BADS: baseline ADAScog
##
## ;BMMS: baseline MMSE
##
## ;;PSTS: 1=normal, 2=MCI, 2=AD
##
## ;APOF: ApoE 0=non carrier, 1=hetero, 2=homo-carrier
##
## ;FHAD: family history of AD, 1=either M or F is AD, 0=no, or NA
## $DATA ../cleaned_800_13_dur36_exp_iiv_0_omg0.csv IGNORE=@
## $ABB COMRES=3
## $PRED
##
## ;Baseline model
## IF (NEWIND.LE.0) THEN
##   LN2=LOG(2)
## ENDIF
##
## INTERCEPT = THETA(1)
## BSLP      = THETA(2)
## TVBSL     = INTERCEPT + BSLP * BMMS
## BASELINE  = TVBSL + ETA(1)
##
## ;----PLACEBO MODEL-----
## PMAX    = THETA(5)+ETA(2)
## HL      = THETA(6)
## PLACEBO = PMAX*(1-EXP(-LOG(2)/HL*TIME))
##
## ;----DRUG MODEL-----
## DRUG = THETA(7)*TIME
##
## IF(MIXNUM.EQ.1) ADASCOG = BASELINE + PLACEBO + DRUG
## IF(MIXNUM.EQ.2) ADASCOG = BASELINE + PLACEBO
## IF(TIME.EQ.0) ADASCOG = BASELINE + PLACEBO
## EST=MIXEST
##
## F = ADASCOG
## ADD = THETA(3)
## W = ADD
## IPRED = F
## Y = F + W * EPS(1)
##
## IRES = DV – IPRED
## IWRES = IRES/W
##
## $CONTR DATA=(ARM)
##
## $MIX
## NSPOP = 2; ARM = 0: placebo ; ARM = 1: treatment
## PMIX1 = THETA(4)
## P(1) = PMIX1
## P(2) = 1 - P(1)
##
## $THETA (0, 62.1334) ; PRM INTER 1
## $THETA (-1.88) ; PRM BSLP 2
## $THETA (0,3) ; PRM EPS 3
## $THETA (0,0.75,1) FIX; PRM MIX 4
## $THETA 21.2001 ; PRM PMAX 5
## $THETA (0,85.3398,100) ; PRM HL 6
## $THETA 1 ; PRM DRUG 7
##
## $OMEGA BLOCK(2)
##    15.386 ; PRM OMBASE 1
##    70.2726 1277.17 ; PRM OMPMAX 2
##
## $SIGMA 1 FIXED
##
## $ESTIMATION MAXEVAL=9999 METHOD=1 NOABORT NOTHETABOUNDTEST NOOMEGABOUNDTEST
~~~

#### 8.1.4 IMA full model

~~~
## $  PROBLEM ADAS-cog
## $  INPUT ID TIME DV BMMS ARM ;$INPUT C ID DROP TIME DV MMSE BADS BMMS DROP DROP DROP
##
## ;PSTS AGE SEX DROP EDU APOE APOF FHAD ABET DROP MDV INVF ARM
##
## ;ADAS=DV
##
## ;TIME in month
##
## ;BADS: baseline ADAScog
##
## ;BMMS: baseline MMSE
##
## ;;PSTS: 1=normal, 2=MCI, 2=AD
##
## ;APOF: ApoE 0=non carrier, 1=hetero, 2=homo-carrier
##
## ;FHAD: family history of AD, 1=either M or F is AD, 0=no, or NA
## $DATA ../cleaned_800_13_dur36_exp_iiv_0_omg0.csv IGNORE=@
## $ABBREVIATED COMRES=3
## $PRED
##
## ;Baseline model
## IF (NEWIND.LE.0) THEN
##   LN2=LOG(2)
## ENDIF
##
## INTERCEPT = THETA(1)
## BSLP      = THETA(2)
## TVBSL     = INTERCEPT + BSLP * BMMS
## BASELINE  = TVBSL + ETA(1)
##
## ;----PLACEBO MODEL-----
## PMAX    = THETA(5)+ETA(2)
## HL      = THETA(6)
## PLACEBO = PMAX*(1-EXP(-LOG(2)/HL*TIME))
##
## ;----DRUG MODEL-----
## DRUG = THETA(7)*TIME
##
## IF(MIXNUM.EQ.1) ADASCOG = BASELINE + PLACEBO + DRUG
## IF(MIXNUM.EQ.2) ADASCOG = BASELINE + PLACEBO
## IF(TIME.EQ.0) ADASCOG   = BASELINE + PLACEBO
## EST=MIXEST
##
## F = ADASCOG
## ADD = THETA(3)
## W = ADD
## IPRED = F
## Y = F + W * EPS(1)
##
## IRES = DV – IPRED
## IWRES = IRES/W
##
## $CONTR   DATA=(ARM)
## $MIX
## NSPOP = 2; ARM = 0: placebo ; ARM = 1: treatment
## PMIX1 = THETA(4)
## P(1) = ((PMIX1+0.5)*ARM+(0.5-3*PMIX1)*(1-ARM))*1.5
## P(2) = 1 - P(1)
##
## $THETA (0,52.7227) ; PRM INTER 1
## $THETA -1.53645 ; PRM BSLP 2
## $THETA (0,2.81556) ; PRM EPS 3
## $THETA (-0.0555,0.0000000001,0.1666) ; PRM MIX 4
## $THETA 10.05 ; PRM PMAX 5
## $THETA (0,61.2471,100) ; PRM HL 6
## $THETA 0.074134 ; PRM DRUG 7
## $OMEGA BLOCK(2)
##    15.2454 ; PRM OMBASE 1
##    51.746 705.833 ; PRM OMPMAX 2
## $SIGMA 1 FIX
## $ESTIMATION MAXEVAL=9999 METHOD=1 NOABORT NOTHETABOUNDTEST
##       NOOMEGABOUNDTEST
~~~

#### 8.1.5 sIMA base model

~~~
## $  PROBLEM ADAS-cog
## $  INPUT ID TIME DV BMMS ARM ;$INPUT C ID DROP TIME DV MMSE BADS BMMS DROP DROP DROP
##
## ;PSTS AGE SEX DROP EDU APOE APOF FHAD ABET DROP MDV INVF ARM
##
## ;ADAS=DV
##
## ;TIME in month
##
## ;BADS: baseline ADAScog
##
## ;BMMS: baseline MMSE
##
## ;;PSTS: 1=normal, 2=MCI, 2=AD
##
## ;APOF: ApoE 0=non carrier, 1=hetero, 2=homo-carrier
##
## ;FHAD: family history of AD, 1=either M or F is AD, 0=no, or NA
## $DATA ../cleaned_800_13_dur36_exp_iiv_0_omg0.csv IGNORE=@
## $ABBREVIATED COMRES=3
## $PRED
##
## ;Baseline model
## IF (NEWIND.LE.0) THEN
##   LN2=LOG(2)
## ENDIF
##
## INTERCEPT = THETA(1)
## BSLP      = THETA(2)
## TVBSL     = INTERCEPT + BSLP * BMMS
## BASELINE  = TVBSL + ETA(1)
##
## ;----PLACEBO MODEL-----
## PMAX    = THETA(5)+ETA(2)
## HL      = THETA(6)
## PLACEBO = PMAX*(1-EXP(-LOG(2)/HL*TIME))
##
## ;----DRUG MODEL-----
## DRUG = THETA(7)*TIME
##
## IF(MIXNUM.EQ.1) ADASCOG = BASELINE + PLACEBO + DRUG
## IF(MIXNUM.EQ.2) ADASCOG = BASELINE + PLACEBO
## IF(TIME.EQ.0) ADASCOG = BASELINE + PLACEBO
## EST=MIXEST
##
## F = ADASCOG
## ADD = THETA(3)
## W = ADD
## IPRED = F
## Y = F + W * EPS(1)
##
## IRES = DV - IPRED
## IWRES = IRES/W
##
## $CONTR DATA=(ARM)
## $MIX
## NSPOP = 2; ARM = 0: placebo ; ARM = 1: treatment
## PMIX1 = THETA(4)
## P(1) = PMIX1
## P(2) = 1 - P(1)
##
## $THETA (0,60.4528849444118) ; PRM INTER 1
## $THETA -2.02344626270837 ; PRM BSLP 2
## $THETA (0,3.07870473740152) ; PRM EPS 3
## $THETA (0,0.683782864919185,1) ; PRM MIX 4
## $THETA (0,22.4102203457104) ; PRM PMAX 5
## $THETA (0,81.8357793093543,100) ; PRM HL 6
## $THETA 1.04785582235444 ; PRM DRUG 7
## $OMEGA BLOCK(2)
## 16.8370806842093 ; PRM OMBASE 1
## 63.538425910038 1336.26519236903 ; PRM OMPMAX 2
## $SIGMA 1 FIX
## $ESTIMATION MAXEVAL=9999 METHOD=1 NOABORT NOTHETABOUNDTEST
##             NOOMEGABOUNDTEST
~~~

#### 8.1.6 sIMA full model

~~~
## $  PROBLEM ADAS-cog
## $  INPUT ID TIME DV BMMS ARM ;$INPUT C ID DROP TIME DV MMSE BADS BMMS DROP DROP DROP
##
## ;PSTS AGE SEX DROP EDU APOE APOF FHAD ABET DROP MDV INVF ARM
##
## ;ADAS=DV
##
## ;TIME in month
##
## ;BADS: baseline ADAScog
##
## ;BMMS: baseline MMSE
##
## ;;PSTS: 1=normal, 2=MCI, 2=AD
##
## ;APOF: ApoE 0=non carrier, 1=hetero, 2=homo-carrier
##
## ;FHAD: family history of AD, 1=either M or F is AD, 0=no, or NA
## $DATA ../cleaned_800_13_dur36_exp_iiv_0_omg0.csv IGNORE=@
## $ABBREVIATED COMRES=3
## $PRED
##
## ;Baseline model
## IF (NEWIND.LE.0) THEN
##   LN2=LOG(2)
## ENDIF
##
## INTERCEPT = THETA(1)
## BSLP      = THETA(2)
## TVBSL     = INTERCEPT + BSLP * BMMS
## BASELINE  = TVBSL + ETA(1)
##
## ;----PLACEBO MODEL-----
## PMAX    = THETA(5)+ETA(2)
## HL      = THETA(6)
## PLACEBO = PMAX*(1-EXP(-LOG(2)/HL*TIME))
##
## ;----DRUG MODEL-----
## DRUG = THETA(7)*TIME
##
## IF(MIXNUM.EQ.1) ADASCOG = BASELINE + PLACEBO + DRUG
## IF(MIXNUM.EQ.2) ADASCOG = BASELINE + PLACEBO
## IF(TIME.EQ.0) ADASCOG   = BASELINE + PLACEBO
## EST=MIXEST
##
## F = ADASCOG
## ADD = THETA(3)
## W = ADD
## IPRED = F
## Y = F + W * EPS(1)
##
## IRES = DV - IPRED
## IWRES = IRES/W
##
## $CONTR DATA=(ARM)
## $MIX
## NSPOP = 2; ARM = 0: placebo ; ARM = 1: treatment
## PMIX1 = THETA(4)
## PMIX2 = THETA(8)
## P(1) = PMIX1*ARM+PMIX2*(1-ARM)
## P(2) = 1 - P(1)
##
## $THETA (0,56.4592) ; PRM INTER 1
## $THETA -1.67667 ; PRM BSLP 2
## $THETA (0,2.79534) ; PRM EPS 3
## $THETA (0,0.0762061,1) ; PRM MIX 4
## $THETA (0,5.25876) ; PRM PMAX 5
## $THETA (0,34.2199,100) ; PRM HL 6
## $THETA 0.934977 ; PRM DRUG 7
## $THETA (0,0.0762061,1) ; PRM MIX2 8
## $OMEGA BLOCK(2)
##   15.1453 ; PRM OMBASE 1
##   16.0783 67.5087 ; PRM OMPMAX 2
## $SIGMA 1 FIX
## $ESTIMATION MAXEVAL=9999 METHOD=1 NOABORT NOTHETABOUNDTEST
##               NOOMEGABOUNDTEST
~~~

#### 8.1.7 IMA-DR base model

~~~
## $PROBLEM ADAS-cog
## $INPUT ID TIME DV BMMS ARM TRT DOSE RPDOSE
## $ABBREVIATED PROTECT
## ;ADAS=DV
##
## ;TIME in month
##
## ;BADS: baseline ADAScog
##
## ;BMMS: baseline MMSE
##
## ;PSTS: 1=normal, 2=MCI, 2=AD
##
## ;APOF: ApoE 0=non carrier, 1=hetero, 2=homo-carrier
##
## ;FHAD: family history of AD, 1=either M or F is AD, 0=no, or NA
## $ABBREVIATED COMRES=3
## $DATA ../cleaned_800_13_dur36_exp_iiv_0512_omg0.5625_1.csv IGNORE=@
## $PRED
##
## ;Baseline model
## IF (NEWIND.LE.0) THEN
##   LN2=LOG(2)
## ENDIF
##
## INTERCEPT = THETA(1)
## BSLP      = THETA(2)
## TVBSL     = INTERCEPT + BSLP * BMMS
## BASELINE  = TVBSL + ETA(1)
##
## ;----PLACEBO MODEL-----
## PMAX    = THETA(5)+ETA(2)
## HL      = THETA(6)
## PLACEBO = PMAX*(1-EXP(-LOG(2)/HL*TIME))
##
## ;----DRUG MODEL-----
## EXTRA = THETA(7)
##
## IF(MIXNUM.EQ.1) ADASCOG = BASELINE + PLACEBO + EXTRA
## IF(MIXNUM.EQ.2) ADASCOG = BASELINE + PLACEBO
## IF(TIME.EQ.0) ADASCOG = BASELINE
## EST=MIXEST
##
## F = ADASCOG
## ADD = THETA(3)
## W = ADD
## IPRED = F
## Y = F + W * EPS(1)
##
## IRES = DV - IPRED
## IWRES = IRES/W
##
## $CONTR DATA=(TRT)
## $MIX
## NSPOP = 2; ARM = 0: placebo ; ARM = 1: treatment
## X=1/4
## DMIX = THETA(4)
## P(1) = 2*(1-X)*((0.5+DMIX)*TRT+(0.5+(X-1)/X*DMIX)*(1-TRT))
## P(2) = 1 - P(1)
##
## $THETA (0,52.5418) ; PRM INTER 1
## $THETA -1.51931 ; PRM BSLP 2
## $THETA (0,2.85665) ; PRM EPS 3
## $THETA (-0.055,0,.166) FIX ; PRM DMIX 4
## $THETA 11.5139 ; PRM PMAX 5
## $THETA (0,33.9328,100) ; PRM HL 6
## $THETA 1.167 ; PRM EXTRA 7
## $OMEGA BLOCK(2)
##    14.9084 ; PRM OMBASE 1
##    32.189 269.563 ; PRM OMPMAX 2
## $SIGMA 1 FIX
## $ESTIMATION MAXEVAL=9999 METHOD=1 NOABORT NOTHETABOUNDTEST
##      NOOMEGABOUNDTEST
~~~

#### 8.1.8 IMA-DR1 full model

~~~
## $PROBLEM  ADAS-cog
## $INPUT    ID TIME DV BMMS ARM TRT DOSE RPDOSE
## $ABBREVIATED PROTECT
## ;ADAS=DV
##
## ;TIME in month
##
## ;BADS: baseline ADAScog
##
## ;BMMS: baseline MMSE
##
## ;PSTS: 1=normal, 2=MCI, 2=AD
##
## ;APOF: ApoE 0=non carrier, 1=hetero, 2=homo-carrier
## ;FHAD: family history of AD, 1=either M or F is AD, 0=no, or NA
## $ABBREVIATED COMRES=3
## $DATA ../cleaned_800_13_dur36_exp_iiv_0512_omg0.5625_1.csv IGNORE=@
## $PRED
##
## ;Baseline model
## IF (NEWIND.LE.0) THEN
##   LN2=LOG(2)
## ENDIF
##
## INTERCEPT = THETA(1)
## BSLP      = THETA(2)
## TVBSL     = INTERCEPT + BSLP * BMMS
## BASELINE  = TVBSL + ETA(1)
##
## ;----PLACEBO MODEL-----
## PMAX    = THETA(5)+ETA(2)
## HL      = THETA(6)
## PLACEBO = PMAX*(1-EXP(-LOG(2)/HL*TIME))
##
## ;----DRUG MODEL-----
## EXTRA = THETA(7)
## EMAX = THETA(8)+ETA(3)
## D50 = THETA(9)+ETA(4)
## RESPOND_DRUG = EMAX*DOSE/(DOSE+D50)
## DRUG = RESPOND_DRUG
##
## IF(MIXNUM.EQ.1) ADASCOG = BASELINE + PLACEBO + DRUG
## IF(MIXNUM.EQ.2) ADASCOG = BASELINE + PLACEBO + EXTRA
## IF(TIME.EQ.0) ADASCOG   = BASELINE
## EST=MIXEST
##
## F = ADASCOG
## ADD = THETA(3)
## W = ADD
## IPRED = F
## Y = F + W * EPS(1)
##
## IRES = DV - IPRED
## IWRES = IRES/W
##
## $CONTR DATA=(TRT)
## $MIX
## NSPOP = 2; ARM = 0: placebo ; ARM = 1: treatment
## X=1/4
## DMIX = THETA(4)
## P(1) = 2*(1-X)*((0.5+DMIX)*TRT+(0.5+(X-1)/X*DMIX)*(1-TRT))
## P(2) = 1 - P(1)
##
## $THETA (0,52.5418) ; PRM INTER 1
## $THETA -1.51931 ; PRM BSLP 2
## $THETA (0,2.85665) ; PRM EPS 3
## $THETA (-0.055,0.01,.166) ; PRM DMIX 4
## $THETA 11.5139 ; PRM PMAX 5
## $THETA (0,33.9328,100) ; PRM HL 6
## $THETA 0.01 ; PRM EXTRA 7
## $THETA 2.5 ; PRM EMAX 8
## $THETA (0,1) ; PRM D50 9
## $OMEGA BLOCK(2)
##    14.9084 ; PRM OMBASE 1
##    32.189 269.563 ; PRM OMPMAX 2
## $OMEGA 0.5625 ; PRM OMEMAX 3
## $OMEGA 0 FIX ; PRM OMD50 4
## $SIGMA 1 FIX
## $ESTIMATION MAXEVAL=9999 METHOD=1 NOABORT NOTHETABOUNDTEST
##       NOOMEGABOUNDTEST
~~~

#### 8.1.9 IMA-DR2 full model

~~~
## $PROBLEM  ADAS-cog
## $INPUT    ID TIME DV BMMS ARM TRT DOSE RPDOSE
## $ABBREVIATED PROTECT
## ;ADAS=DV
##
## ;TIME in month
##
## ;BADS: baseline ADAScog
##
## ;BMMS: baseline MMSE
##
## ;PSTS: 1=normal, 2=MCI, 2=AD
##
## ;APOF: ApoE 0=non carrier, 1=hetero, 2=homo-carrier
##
## ;FHAD: family history of AD, 1=either M or F is AD, 0=no, or NA
## $ABBREVIATED COMRES=3
## $DATA ../cleaned_800_13_dur36_exp_iiv_0512_omg0.5625_1.csv IGNORE=@
## $PRED
##
## ;Baseline model
## IF (NEWIND.LE.0) THEN
##   LN2=LOG(2)
## ENDIF
##
## INTERCEPT = THETA(1)
## BSLP      = THETA(2)
## TVBSL     = INTERCEPT + BSLP * BMMS
## BASELINE  = TVBSL + ETA(1)
##
## ;----PLACEBO MODEL-----
## PMAX    = THETA(5)+ETA(2)
## HL      = THETA(6)
## PLACEBO = PMAX*(1-EXP(-LOG(2)/HL*TIME))
##
## ;----DRUG MODEL-----
## EXTRA = THETA(7)
## EMAX = THETA(8)+ETA(3)
## D50 = THETA(9)+ETA(4)
## RESPOND_DRUG = EMAX*DOSE/(DOSE+D50)
## DRUG = RESPOND_DRUG
##
## IF(MIXNUM.EQ.1) ADASCOG = BASELINE + PLACEBO + DRUG + EXTRA
## IF(MIXNUM.EQ.2) ADASCOG = BASELINE + PLACEBO + EXTRA
## IF(TIME.EQ.0) ADASCOG = BASELINE
## EST=MIXEST
##
## F = ADASCOG
## ADD = THETA(3)
## W = ADD
## IPRED = F
## Y = F + W * EPS(1)
##
## IRES = DV - IPRED
## IWRES = IRES/W
##
## $CONTR DATA=(TRT)
## $MIX
## NSPOP = 2; ARM = 0: placebo ; ARM = 1: treatment
## X=1/4
## DMIX = THETA(4)
## P(1) = 2*(1-X)*((0.5+DMIX)*TRT+(0.5+(X-1)/X*DMIX)*(1-TRT))
## P(2) = 1 - P(1)
##
## $THETA (0,52.5418) ; PRM INTER 1
## $THETA -1.51931 ; PRM BSLP 2
## $THETA (0,2.85665) ; PRM EPS 3
## $THETA (-0.055,0.01,.166) ; PRM DMIX 4
## $THETA 11.5139 ; PRM PMAX 5
## $THETA (0,33.9328,100) ; PRM HL 6
## $THETA 0.01 ; PRM EXTRA 7
## $THETA 2.5 ; PRM EMAX 8
## $THETA (0,1) ; PRM D50 9
## $OMEGA BLOCK(2)
##   14.9084 ; PRM OMBASE 1
##   32.189 269.563 ; PRM OMPMAX 2
## $OMEGA 0.5625 ; PRM OMEMAX 3
## $OMEGA 0 FIX ; PRM OMD50 4
## $SIGMA 1 FIX
## $ESTIMATION MAXEVAL=9999 METHOD=1 NOABORT NOTHETABOUNDTEST
##       NOOMEGABOUNDTEST
~~~

### 8.2 Mixture proportion domain depending on the allocation ratio

**Figure 4:**
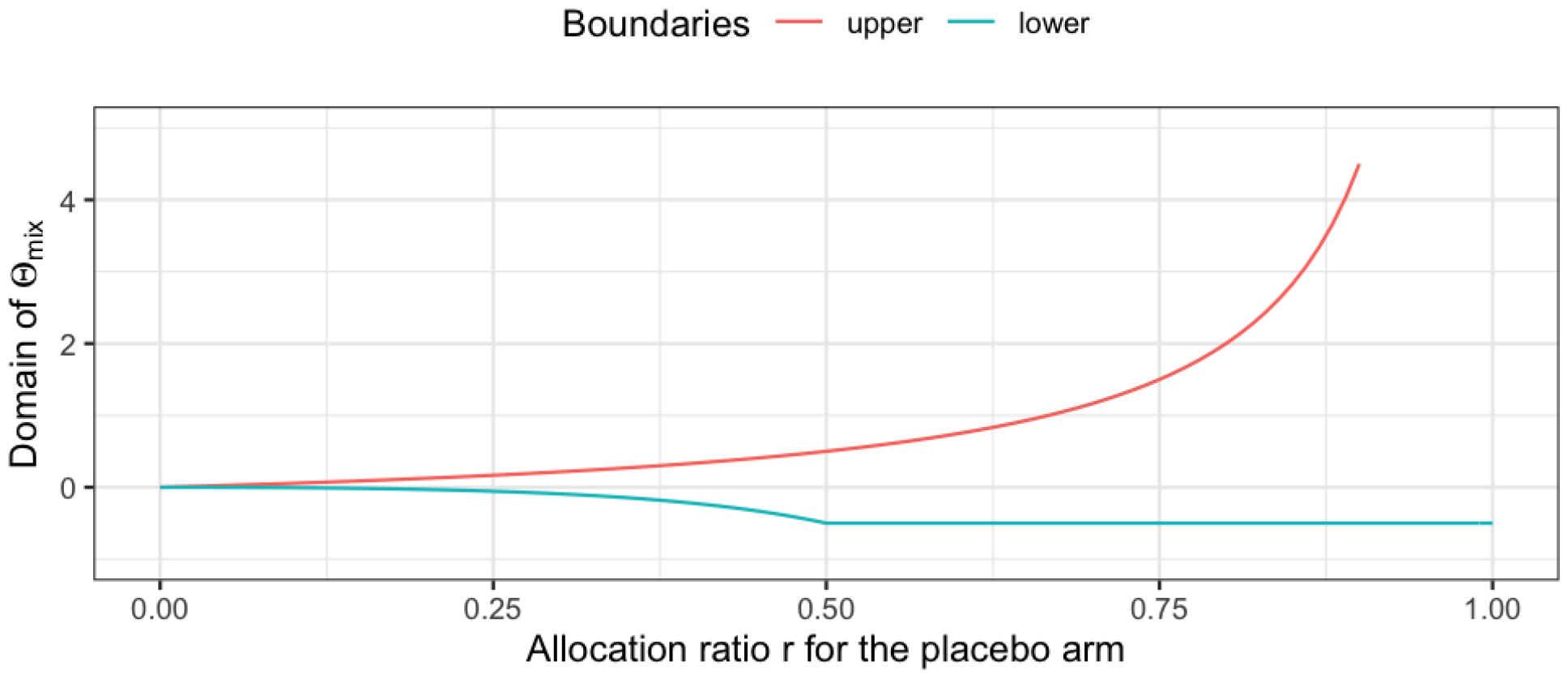
Illustration of the mixture proportion domain depending on the placebo allocation ratio.

### 8.3 Unbalanced design runs details

The minimization status of the models fitted to assess the type I error rate for the unbalanced design for the three placebo allocation rates are presented in Figure 5 for STD, in Figure 6 for IMA, and in Figure 7 for sIMA. For each scenario 1000 base models and 1000 full models were fitted.

**Figure 5:**
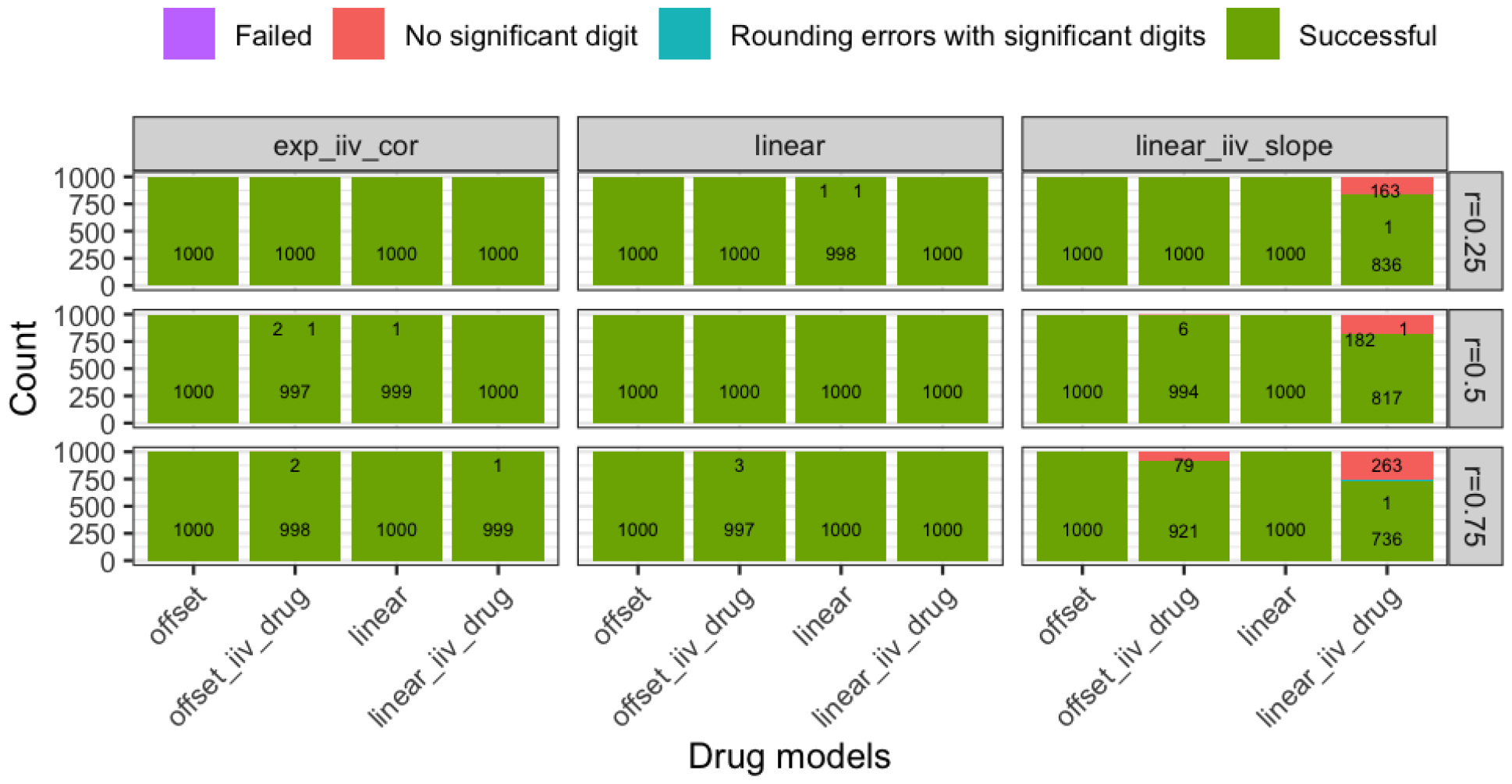
Minimization status for the models fitted on the placebo data to assess the type I error rate for the STD approach.

**Figure 6:**
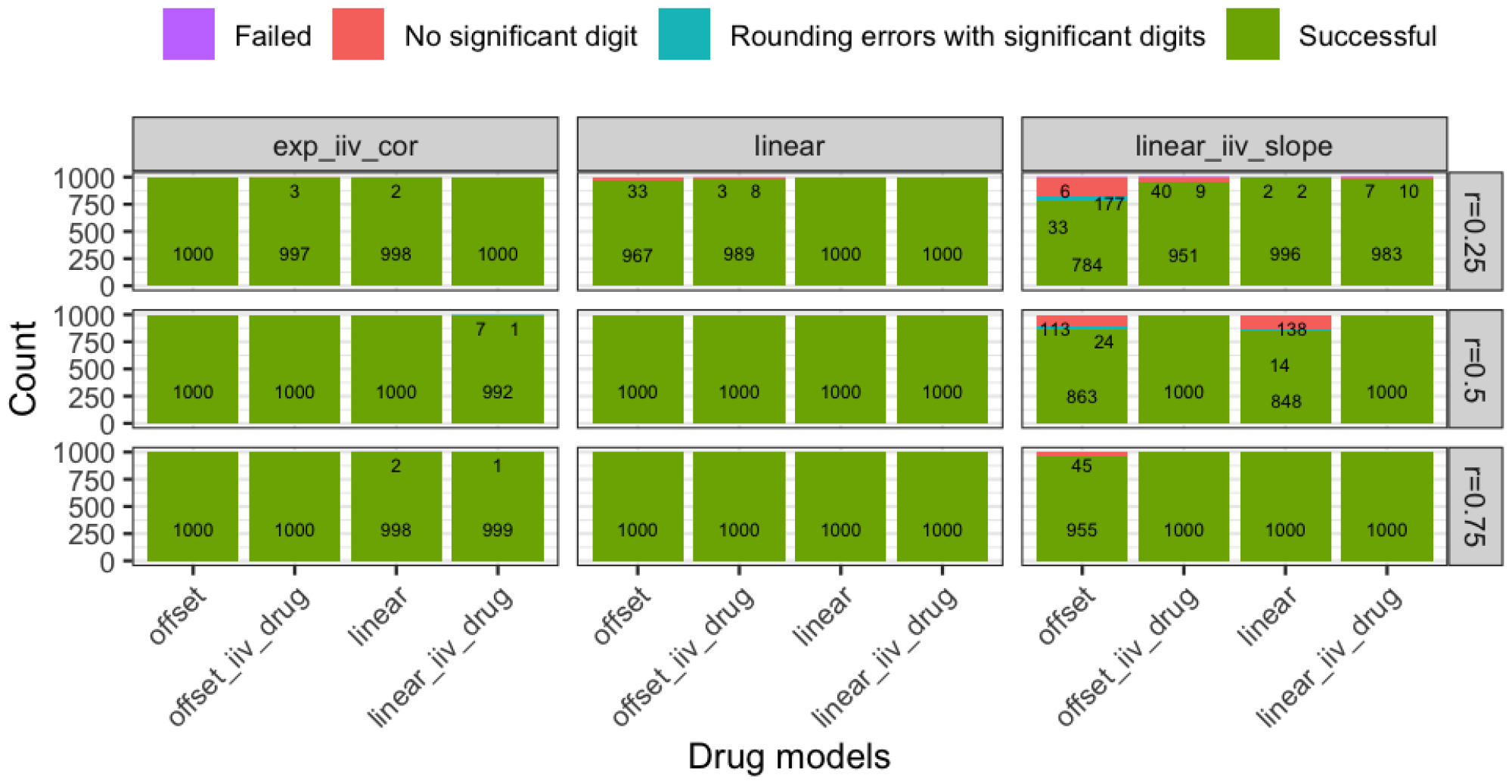
Minimization status for the models fitted on the placebo data to assess the type I error rate for the IMA approach.

**Figure 7:**
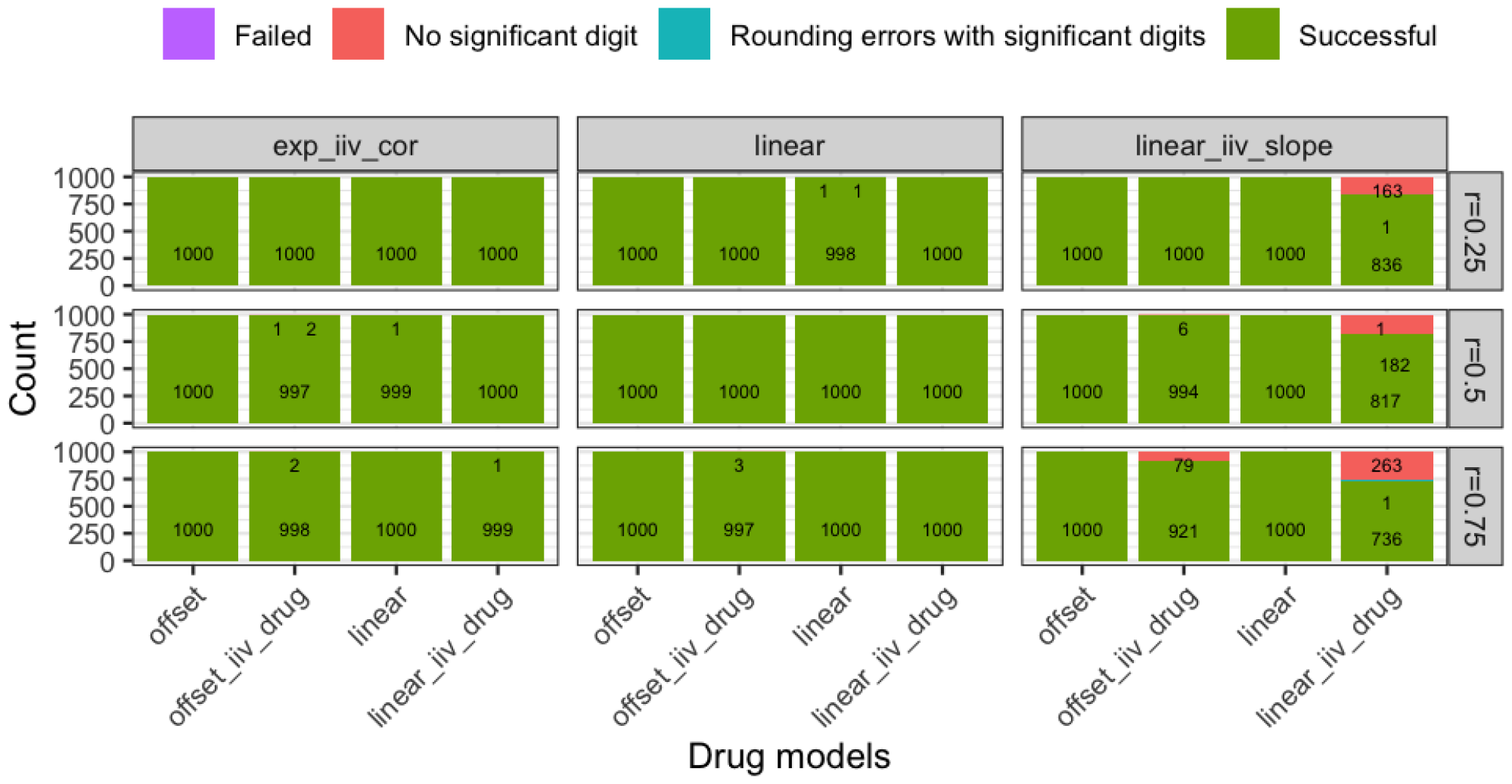
Minimization status for the models fitted on the placebo data to assess the type I error rate for the sIMA approach.

### 8.4 Dose-response runs details

The minimization status of the models fitted to assess the bias in the drug effect estimates for data modified by the addition of various dose-response scenarios is presented in Figure 8. For each scenario 1000 base models and 1000 full models were fitted. Ths mixture proportion estimated for IMA-DR1 and IMA-DR2 is presented in Figure 8 for all the simulated scenarios.

**Figure 8:**
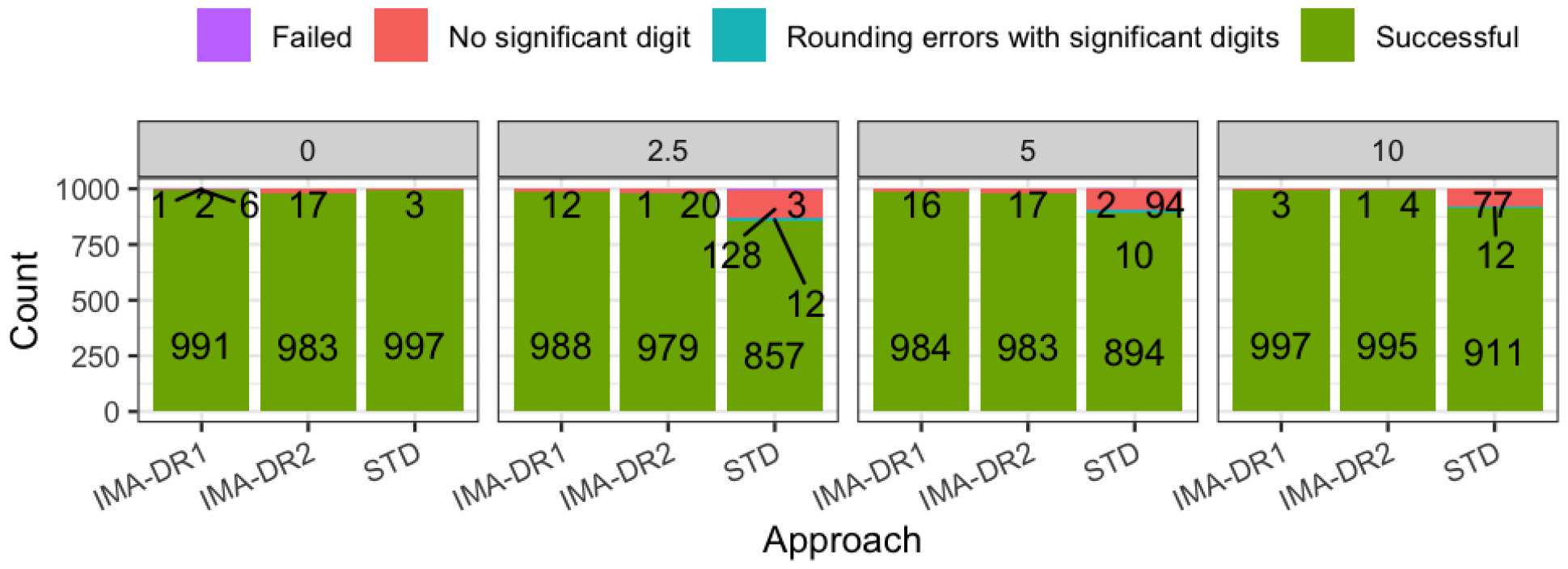
Minimization status for the models fitted on the placebo data modified by the addition of dose-response to assess the bias in the drug estimates for STD, IMA-DR1, and IMA-DR2.

**Figure 9:**
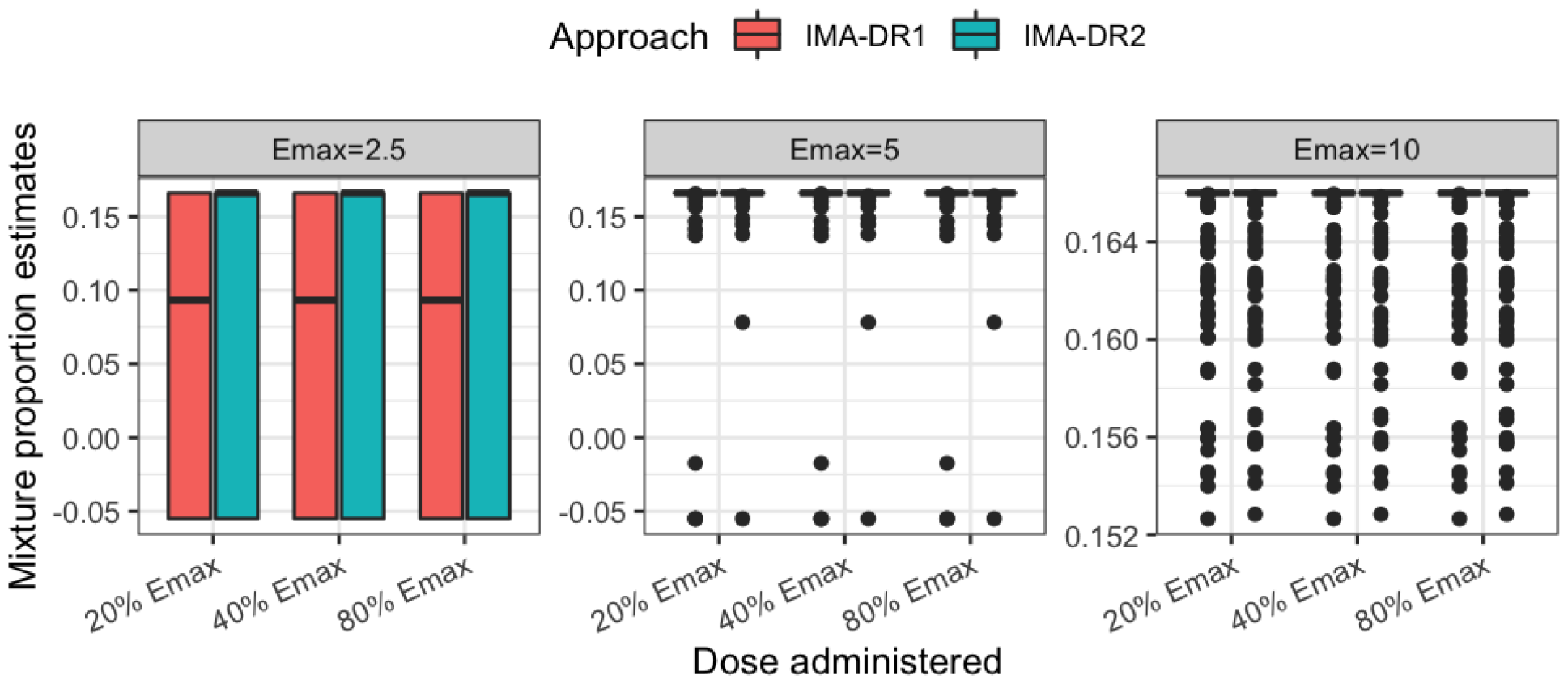
Mixture proportion estimate for each of the dose-response scenarios simulated, using the placebo data modified by the addition of a dose-response (N=1000).

## References

1. Health UD of, Services H, Food, et al (2004) Innovation or stagnation. Challenge and opportunity on the critical path to new medical products

2. Bhattaram VA, Booth BP, Ramchandani RP, et al (2005) Impact of pharmacometrics on drug approval and labeling decisions: A survey of 42 new drug applications. The AAPS journal 7:E503–E512

3. Powell J, Gobburu J (2007) Pharmacometrics at FDA: Evolution and impact on decisions. Clinical Pharmacology & Therapeutics 82:97–102

4. Miller R, Ewy W, Corrigan BW, et al (2005) How modeling and simulation have enhanced decision making in new drug development. Journal of pharmacokinetics and pharmacodynamics 32:185–197

5. Lalonde R, Kowalski K, Hutmacher M, et al (2007) Model-based drug development. Clinical Pharmacology & Therapeutics 82:21–32

6. Chasseloup E, Tessier A, Karlsson MO (2021) Assessing treatment effects with pharmacometric models: A new method that addresses problems with standard assessments. The AAPS journal 23:1–8

7. Carlsson KC, Savi RM, Hooker AC, Karlsson MO (2009). The AAPS journal 11:148–154

8. Beal SL, Sheiner LB, Boeckmann AJ, and Bauer RJ (eds) (1989-2018) NONMEM 7.4 users guides. ICON plc, Gaithersburg, MD

9. Keizer RJ HA Karlsson MO (2013) Modeling and simulation workbench for NONMEM: Tutorial on pirana, PsN, and xpose. CPT Pharmacometrics Syst Pharmacol 2:e50

10. R Core Team (2019) R: A language and environment for statistical computing. R Foundation for Statistical Computing, Vienna, Austria

11. Ito K, Corrigan B, Zhao Q, et al (2011) Disease progression model for cognitive deterioration from alzheimer’s disease neuroimaging initiative database. Alzheimer’s & Dementia 7:151–160

